# Recurrent chromosomal imbalances provide selective advantage to human embryonic stem cells under enhanced replicative stress conditions

**DOI:** 10.1101/2020.11.06.360685

**Authors:** Liselot M. Mus, Stéphane Van Haver, Mina Popovic, Wim Trypsteen, Steve Lefever, Nadja Zeltner, Yudelca Ogando, Eva Z. Jacobs, Geertrui Denecker, Ellen Sanders, Christophe Van Neste, Suzanne Vanhauwaert, Bieke Decaesteker, Dieter Deforce, Filip Van Nieuwerburgh, Pieter Mestdagh, Jo Vandesompele, Björn Menten, Katleen De Preter, Lorenz Studer, Björn Heindryckx, Kaat Durinck, Stephen Roberts, Frank Speleman

**Affiliations:** Department of Biomolecular Medicine, Ghent University, Ghent, Belgium; Cancer Research Institute Ghent (CRIG), Ghent, Belgium; Ghent-Fertility and Stem Cell Team (G-FaST), Department for Reproductive Medicine, Ghent University Hospital, Ghent, Belgium; Center for Molecular Medicine, Department of Biochemistry & Molecular Biology and Department of Cellular Biology, University of Georgia, Athens, GA, USA; Department of Pediatrics, Memorial Sloan Kettering Cancer Center, New York, USA; Laboratory of Pharmaceutical Biotechnology, Ghent University, Ghent, Belgium; The Center for Stem Cell Biology, Sloan Kettering Institute, New York, USA; Developmental Biology Program, Sloan Kettering Institute, New York, USA

**Keywords:** hESC, replicative stress, copy number variations

## Abstract

Human embryonic stem cells (hESCs) and embryonal tumors share a number of common features including a compromised G1/S checkpoint. Consequently, these rapidly dividing hESCs and cancer cells undergo elevated levels of replicative stress which is known to induce genomic instability causing chromosomal imbalances. In this context, it is of interest that long-term *in vitro* cultured hESCs exhibit a remarkable high incidence of segmental DNA copy number gains, some of which are also highly recurrent in certain malignancies such as 17q gain (17q+). The selective advantage of DNA copy number changes in these cells has been attributed to several underlying processes including enhanced proliferation. We hypothesized that these recurrent chromosomal imbalances become rapidly embedded in the cultured hESCs through a replicative stress driven Darwinian selection process. To this end, we compared the effect of hydroxyurea induced replicative stress versus normal growth conditions in an equally-mixed cell population of isogenic euploid and 17q+ hESCs. We could show that 17q+ hESCs rapidly overtook normal hESCs. Our data suggest that recurrent chromosomal segmental gains provide a proliferative advantage to hESCs under increased replicative stress, a process that may also explain the highly recurrent nature of certain imbalances in cancer.

## INTRODUCTION

Human embryonic stem cells (hESCs) can be maintained indefinitely as self-renewing cell populations, while retaining the capacity to differentiate into all cell lineages ^1^. During normal embryonal development, pluripotent cells of the inner cell mass, equivalent to hESC, exist only transiently. Highly proliferative mouse (m)ESCs are shown to exhibit a short G1-phase due to a compromised G1/S checkpoint. This causes endogenous replicative stress and subsequent DNA damage ^2^, which may be further enhanced under *in vitro* culture conditions. This could explain the frequent occurrence of chromosomal imbalances in long term cultured hESC lines. Further, the highly recurrent nature of chromosome 12 and 17q gains (17q+) suggests a strong selective pressure towards outgrowth of these subclones ^3–5^. The functional relevance of these copy number gains is further supported by the finding of a trisomy for chromosome 11, syntenic to human chromosome 17, in almost 20% of mESCs lines ^3^. Of further interest, Zhang *et al.* reported an increased proliferation rate, colony formation efficiency and teratoma formation for aneuploid mESC lines (including trisomy 11)^6^. This is of interest given that 17q gains and amplifications are a recurrent finding in cancers including embryonic malignancies such as neuroblastoma and medulloblastoma. In neuroblastoma, we recently reported an elevated ESC-derived stemness gene signature activity in ultra-high-risk patients ^7^. In view of the above, we investigated the selective growth advantage of segmental chromosomal aberrations (with primary focus on 17q+) in hESCs as this may also provide insight into the functional significance of recurrent 17q+ in cancer cells. We hypothesized that acquired recurrent DNA copy number imbalances (in particular 17q+) in hESC cell lines, reflect an adaptive process to protect cells from excessive replicative stress. To test this, we monitored selective growth of isogenic hESCs with 17q+ versus the normal euploid parental line, in normal growth conditions and enhanced replicative stress. We observed rapid overgrowth of 17q+ hESCs over normal hESCs, only in conditions of replicative stress. Of further interest, under these elevated replicative stress conditions, an additional subclone with segmental 13q gain (and varying other smaller imbalances) also emerged over time, suggesting that dosage effects for genes mapping to this chromosomal region also contribute to replicative stress resistance. We propose that the frequently observed segmental chromosomal gains in cultured hESC lines drive an adaptive mechanism to cope with prolonged replicative stress through combined gene dosage effects. Further studies in additional cell lines are warranted to explore the underlying molecular basis of this selective process in more depth, which could also shed light onto similar replicative stress-driven selective processes that shape the highly recurrent DNA copy number landscape in embryonal tumors such as neuroblastoma and medulloblastoma.

## MATERIALS AND METHODS

### Ethical permissions

The hESC line used, UGENT11-60 (C12), is an existing line kindly provided to us by the lab of prof. B. Heindryckx ^8^. The derivation and use of naive hESC was approved by the Ghent University Hospital Ethical Committee (EC2013/822) and by the Belgian Federal Commission for medical and scientific research on embryos *in vitro* (ADV052 UZ Ghent). The study of the role of 17q+ in a hESC-derived MYCN-driven neuroblastoma-model was approved by the Ghent University Hospital Ethical Committee (EC2018/0805). All hESC samples and derivates obtained are deposited according to regulations in the CMGG biobank (FAGG BB190119; EC2019/0392). All experiments were performed in accordance with relevant guidelines and regulations.

### Naive hESC culture

Naive UGENT11-60 hESCs (referred to as C12) were cultured as described previously (Warrier et al., 2017). In brief, naive hESCs were cultured on inactivated (by irradiation) Mouse Embryonic Fibroblasts (MEFs; A34180, Fisher Scientific). MEFs were seeded at a density of 200 000 cells per well of a 6-well plate in MEF media consisting of DMEM 1x (41965-039, Invitrogen) supplemented with 10% fetal calf serum, 1% L-glutamine (25030-025, Invitrogen) and 1% Penicillin/Streptomycin (15140-122, Invitrogen). Cells were allowed to attach and spread prior to use. hESC were subsequently seeded on the attached MEF. Basal naive hESC medium consisted of Knockout DMEM (10829-018, Invitrogen) containing 20% knockout serum replacement (10828-028, Invitrogen), 2mM L-glutamine, 1% Penicillin/Streptomycin, 1% non-essential amino acids (11140-050, Invitrogen), and 0.1mM β-mercapto-ethanol (31350-010, Invitrogen). This basal naive hESC medium is supplemented with 12ng/ml bFGF (100–18B, Peprotech) and 1000U recombinant human LIF (300-05, Sigma) and the small molecules 1μM PD0325901 (13034, Sanbio), 3μM CHIR99021 (1386, Axon Medchem), 10μM Forskolin (F6886, Sigma) and 50ng/ml ascorbic acid (A8960, Sigma) and used as complete media. Naive hESCs were maintained in hypoxic conditions of 5% O2 and 5% CO2 conditions at 37°C. MEF depletion in the hESC culture at passaging was performed by discarding the 0.05% trypsin/EDTA (25300-054, Invitrogen) applied to the culture after an incubation of 5min at room temperature. Next, loosened hESC colonies and remaining MEF were resuspended in complete hESC media. Naive colonies were passaged as single cells at a density of 35 000 cells per well of a 6-well plate every three days and were replated on inactivated MEFs.

### Isolation of monoclonal lines by limited dilution

Due to the observation of chromosomal aberrations upon prolonged culture (> 107 passages), isolation of monoclonal lines by limited dilution was performed to analyse the presence of subclones in the C12-cl28 line at early passage (passage 49). Therefore, hESCs were dissociated into single cells using 0.05% Trypsin/EDTA. To prevent clumping of cells, a cell strainer was used. The cells were diluted to a final concentration of 0.5 cells per 100μl of complete naive medium, supplemented with 10μM Rock inhibitor (S1049, Selleckchem), to reduce the likelihood of having multiple cells per well. 100μl of diluted cells were added to each well of a 96-well plate, coated with inactivated MEFs. After six to eight days, individual colonies were observed and these were picked and further expanded.

### Co-culture experiments

To analyse the subtle differences in growth advantage of 17q gained cells over wild type cells a co-culture experiment was set up. Here, 15 000 cells of either monoclonally grown C12-cl28 or C12-cl32 cell lines (passage 54 and 53 respectively) were mixed, labelled as “mix 50%”, and seeded in two-fold on the MEF feeder layer. The cells were seeded in complete media either supplemented with 150μM hydroxyurea (HU; H8627-1G, Sigma) or equal amounts of PBS (14190-094, Invitrogen) as control condition for a total duration of 60 days. A normal passaging regime, i.e. every three days, was continued and the co-culture was treated as a new cell line. The C12-cl28, C12-cl32 or co-culture were subsequently passaged each time at 30 000 cells/well in control conditions and 40 000 cells/well under HU treatment. Samples for genomic DNA, RNA and FACS analysis were isolated at each passage.

### Morphological assessment

Due to the application of HU in order to induce replicative stress, cells started to obtain an altered morphology. Morphological assessment was done by IncuCyte® live-cell imaging or by using an EVOS phase contrast microscope.

### 3T3 assay

To quantify the proliferation rate over time and several passages, a 3T3-assay was applied as previously described (Todaro and Green, 1963). In brief, at each passage colonies were disrupted by applying trypsin, as described earlier, and a single cell suspension was obtained. Cells of C12-cl28, C12-cl32 or co-culture, either maintained in control conditions (PBS) or induced replicative stress (HU), were plated in 6-well plates at the densities described during normal culture. Three days later, the total number of cells was counted again at passaging and cells were re-plated. The cumulative increase in cell number was calculated according to the formula LOG (Nf/Ni)/LOG2, where Ni and Nf are the initial and final numbers of cells plated and counted after three days respectively. Error bars in figures represent SD after error propagation. For statistical testing, each growth curve was modelled using a linear model, each model explained at least 97% of the observed variance. Next, models were tested for equal slope using lstrends (lsmeans package) at 1% significance (Lenth, 2016).

### Genomic DNA extraction

Cell pellets were dissolved in 150μl PBS. Genomic DNA was subsequently extracted by QIAcube (Qiagen) using the QiaAmp Blood mini kit (250, Qiagen) consisting of AL, AE, AW1 and AW2 buffers, Qiagen protease, QIAmp mini spin columns and collection tubes. DNA extraction was conducted according to the manufacturer’s instructions. DNA concentrations were measured by using the Dropquant software of the Dropsense. Samples were diluted appropriately and used for targeted sequencing, sWGS and ddPCR.

### Shallow-Whole Genome Sequencing (sWGS)

For shallow depth WGS (sWGS), 312ng of extracted gDNA was first diluted in TE buffer (12090-015, Invitrogen) to a total volume of 50μl. Next the gDNA was sheared with the help of a Covaris LE220+ focused-ultrasonicator (200bp). The sheared gDNA was subsequently transferred to the Hamilton Star® for automated library preparation where after library concentration measurements were conducted using the Qubit fluorometer HS kit. Finally, the library pools were sequenced using the HiSeq3000. Each genome profile (line view and chromosome view) was manually checked for aberrations. All 115 profiles, co-culture experiment samples were taken at each passage during 60 days, were visualized using the online tool Vivar (http://cmgg.be/vivar/)(Sante et al., 2014) and data is available at GEO (GSE150647).

### Targeted sequencing

Tumor and normal DNA were profiled using the 69-gene MDG SeqCap SOLID panel (https://www.cmgg.be/nl/zorgverlener/labguide/platform-moleculaire-diagnostiek-uz-gent-mdg). This assay involves hybridization of barcoded libraries to custom oligonucleotides (Nimblegen SeqCap) designed to capture all protein-coding exons and select introns of 69 commonly implicated oncogenes, tumor suppressor genes, and members of pathways deemed actionable by targeted therapies. Barcoded sequence libraries were prepared using 100ng genomic DNA with the KAPA HyperPlus Prep Kit (Kapa Biosystems KK8504) and combined in equimolar pools. The captured pools were subsequently sequenced on an Illumina MiSeq as paired-end 150-base pair reads, producing > 300-fold coverage per ROI.

### Digital droplet PCR

As an independent and quantitative validation of the sWGS results, it was chosen to perform a digital droplet PCR (ddPCR) for the chromosomal copy numbers of 17q on the extracted gDNA. All ddPCR reactions were executed according to an in-house optimized and standardized protocol. Since the breakpoint of 17q in C12-cl32 was located at 17q21.31, it was opted to use BRIP1 (17q23.2) as target gene. WWC1 (5q34) was used as reference target. Specific primers and probes were designed using RTPrimerDB (http://www.rtprimerdb.org)(Lefever et al., 2009): as BRIP1 primers CCTGGACACATATCTTTGCT (forward) and GGACAATGAGTCTACACTTGA (reverse) were used; as WWC1 primers GAAGGTACTGAGTCACAAGTC (forward) and AACAACAGTAGGACCCAAAC (reverse) were used and as probes /56-FAM/TGGAAGCAG/ZEN/CAAGTCATCTATCACC/3IABkFQ for BRIP1 and /5-HEX/CGGTTCATC/ZEN/TTCAAACCAACAAGAGC/3IABkFQ for WWC1 were used.

Prior to this study the optimal annealing temperature was investigated for every primer-probe combination. Droplets were generated using the QX100 Droplet Generator (Bio-Rad) from 20 L mixtures containing 5ng of gDNA, 250nM primers, 900nM probes and 10μL ddPCR Supermix for Probes (Bio-Rad). The PCR reaction was executed using a thermocycler with a 10min denaturation step at 95°C followed by repeated cycles consisting of 15s at 95°C and one min at 59°C. After 40 cycles, the products were heated to 98°C for 10min before cooling to room temperature. Results were read out with a QX100Droplet Reader (Bio-Rad). Calling of positive and negative droplets and ratio calculations were performed using the Quantasoft software (Bio-Rad). Quantitative copy number analysis for 17q was performed using generalised linear mixed models (GLMM; http://www.dpcr.ugent.be)(Vynck et al., 2016). Error bars in figures represent 95% CI after error propagation.

### Cell cycle analysis using FACS

For cell cycle analysis, 35 000 cells were seeded per well in duplicates in complete naive media either supplemented with 150μM HU or equal amounts of PBS. Cells were trypsinized and MEFs were depleted as described above. Loosened hESC colonies and remaining MEF were resuspended in complete hESC media and centrifuged at 39g. Next pellets were washed with PBS followed by a hESC specific staining by adding 0.5μl of the TRA-1-60-FITC antibody (560173, BD Pharmingen) per 100 000 cells and incubated for 30min at 4°C in the dark. To avoid overstaining, cells were washed in PBS and were subsequently resuspended in 300μl cold PBS and, while vortexing, 700μl ice-cold ethanol was added dropwise to fix the cells. Following an incubation of the sample for minimum 1h at −20°C, cells were washed in PBS and resuspended in 100μl PBS with RNase A (19101, Qiagen) to a final concentration of 0.25mg/ml. Upon 1h incubation at 37°C, 4μl propidium iodide solution (P4170-25mg, Sigma) were added to a final concentration of 40μg/ml. Samples were loaded on a BioRad S3TM cell sorter and analysed with the Dean-Jett-Fox algorithm for cell-cycle analysis using the ModFit LTTM software package. Error bars in figures represent SD after scaling to control and error propagation. For statistical testing, a Wilcoxon rank sum test was performed at the 5% significance level.

### Transcriptional network analysis

RNA sequencing was carried out on samples collected of C12-cl28 and C12-cl32 under normal culture conditions (PBS) and conditions of replicative stress (150μM HU), for six consecutive passages (passage 1-6) during two independent cultures (n=12 biological replicates per condition, 52 samples in total). Total RNA was isolated using the miRNeasy micro kit (217084, Qiagen) according to manufacturer’s instructions, including on-column DNase treatment. Next, libraries for mRNA sequencing were prepared using the QuantSeq 3’ mRNA FWD kit (Lexogen) prior to sequencing on the Nextseq500 (High output, SR75, Illumina). On average 7.0 ± 1.9M high-quality reads were generated per samples. For each sample, gene-level read counts were generated upon mapping to the Homo sapiens reference genome (GRCh38) using STAR. Counts were normalized using edgeR and differential gene expression profiling was performed using DeSeq2. Gene Set Enrichment Analysis (GSEA)(Subramanian et al., 2005) was performed using the c1 and c2 curated gene collection of MSigDB. RNA sequencing data is available at GEO (GSE150647).

## RESULTS

### Recurrent acquired chromosomal imbalances confer a proliferative advantage to hESCs under conditions of replicative stress

Our lab studies the molecular and functional impact of 17q gain in the context of neuroblastoma, a pediatric tumor of the sympathetic nervous system ^9^. In order to model this further using hESC derived models, we performed copy number analysis in available hESC cell lines and identified chromosome 17q21.31q25.3 gain in the naive hESC line UGENT11-60 (further referred to as C12) at late passage 124 in addition to gains of chromosome 1q25.3q32.1 and chromosome 7 as major aberrations (**Fig. S1A, Table S1**). Subsequent comparison of shallow whole genome sequencing (sWGS) profiles of the subclones following dilution cloning from an earlier passage (passage 49), identified a C12 subclone (C12-cl28) with no DNA copy number variations and a C12 subclone with a large gain of chromosome 17q21.31q25.3, further referred to as 17q+, (C12-cl32; **Fig. 1A & S1B**), as well as a few other small imbalances/deletions (**Table S1**) and without known pathogenic variants of *TP53* or other cancer-related genes as analysed by targeted sequencing. As such, C12-cl32 was used to evaluate the effect of this large chromosome 17q+ on relative proliferative capacity compared to its parental chromosome 17q copy neutral isogenic counterpart. We performed a mixing experiment (1:1 ratio) of the subclones under normal growth conditions versus chronic exposure to hydroxyurea (HU, 150μM), a well-established compound known to induce increased replicative stress levels through inhibition of nucleotide synthesis ^10^. The proportion of both clones was monitored at each passage, over a period of 60 consecutive days through a total of 115 sWGS (**Fig. 1B & S1C**) and 36 digital droplet PCR analyses (ddPCR; **Fig. 1C**). Both sWGS and ddPCR confirmed a 1:1 proportion at the start of the experiment. In the absence of HU, the 1:1 proportion of C12-cl28 and -cl32 cells remained stable or was even slightly reduced for the duration of this experiment. In contrast, in conditions supplemented with HU, the 17q+ containing C12-cl32 cells rapidly overtook the normal cells with 75.1% increase after 21 days and 90.8% after 42 days of culture, suggesting that the presence of 17q+ provides a proliferative advantage, but only under conditions of HU-induced replicative stress (**Fig. 1B**). Of interest, besides the increase in the proportion of hESC with 17q+ during culture under HU, a second apparently pre-existing 17q+ clone with additional 13q21.32q34 gain (82.84%) emerged together with extra smaller imbalances, indicating that in addition to 17q other imbalances were selected (**Table S1)**. Indeed, also under normal growth conditions, the same 13q+/17q+ subclone emerged but at a slower rate compared to the HU-treated culture, both in the C12-cl32 culture (23.46% versus 80.89%) (**Fig. S1C**) and mixing experiment (28.15% versus 82.84%, respectively) (**Fig. 1B**). In the C12-cl32 cells only grown under HU conditions, yet another subclone emerged (partial 2p and 3q gain), suggesting that several 17q+ subclones co-exist with the major 17q+ clone (**Table S1)**.

**Figure 1:**
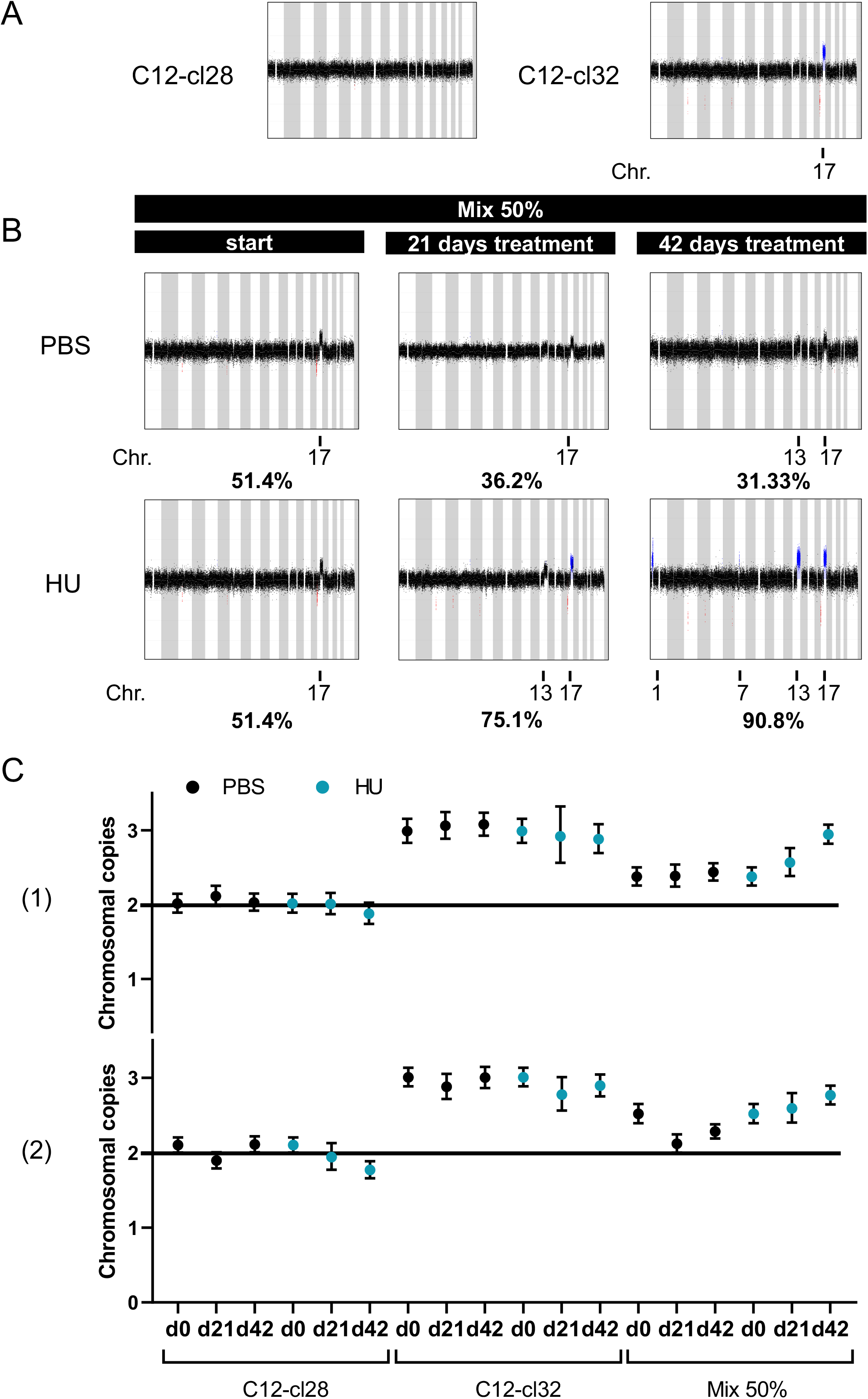
Recurrent acquired chromosomal imbalances confer a proliferative advantage to hESCs under conditions of replicative stress. **[A]** Chromosomal profiles by sWGS of C12-cl28 and C12-cl32. (see also **Table S1**) **[B]** Chromosomal profiles by sWGS of co-culture (mix 50%) over time (days), under control conditions (PBS) and replicative stress (150μM HU). Percentages represent the portion of 17q+ cells in the co-culture (mix 50%; representative of n=2 shown (biological replicates), see also **Fig. S1**) [**C]** Quantitative representation of chromosomal copies of 17q over time (days) by ddPCR of C12-cl28, C12-cl32 and co-culture (mix 50%) under control conditions (PBS) and replicative stress (150μM HU; biological replicates shown as (1) and (2) with 2 technical replicates; copy number with error bars representing the 95% CI)

### Acquired chromosomal imbalances including the common 17q gain attenuate hydroxyurea-induced G1/S-phase arrest and provide a proliferative advantage

To further investigate the presumed effect of chromosome 17q+ on the proliferative capacity, cell cycle analysis was done following HU exposure using flow cytometry analyses during the early phases of the co-culture experiment (passage 3-6) prior to emergence of prominent additional imbalances (**Fig. 2A**). We first excluded the mouse embryonic fibroblast (MEF) feeder cells present in the naive cultures, by sorting based on the human specific pluripotent stem cell surface marker TRA-1-60 (**Fig. S2A**). Subsequently, flow cytometry measurements revealed significant S-phase arrest (p=0.02857) upon HU treatment compared to control for both C12-cl28 and -cl32 cells as well as the co-culture, corresponding to the expected HU-induced replicative stress ^10^ (**Fig. 2A** **& S2B**). Importantly, in 17q+ C12-cl32 cells the proportion of cells with HU-induced increase in S-phase is attenuated with 17% as compared to normal C12-cl28 cells. Comparison of effects of HU-induced increase in S-phase cells versus G2/M-phase cells reveals that while both clones show an increase in S-phase cells with HU treatment, the reduced proportion of cells in S-phase upon HU treatment of C12-cl32 is accompanied with increased G2/M-phase cell population. In contrast, for C12-cl28 cells the proportion of cells in G1- and G2/M-phase remains stable. The corresponding copy number variation profiles confirm the presence of 17q+ in all cells of the C12-cl32 culture and show the emergence of a 13q+/17q+ subclone (20-35% from 10 to 21 days; **Fig. 2B)**.

**Figure 2:**
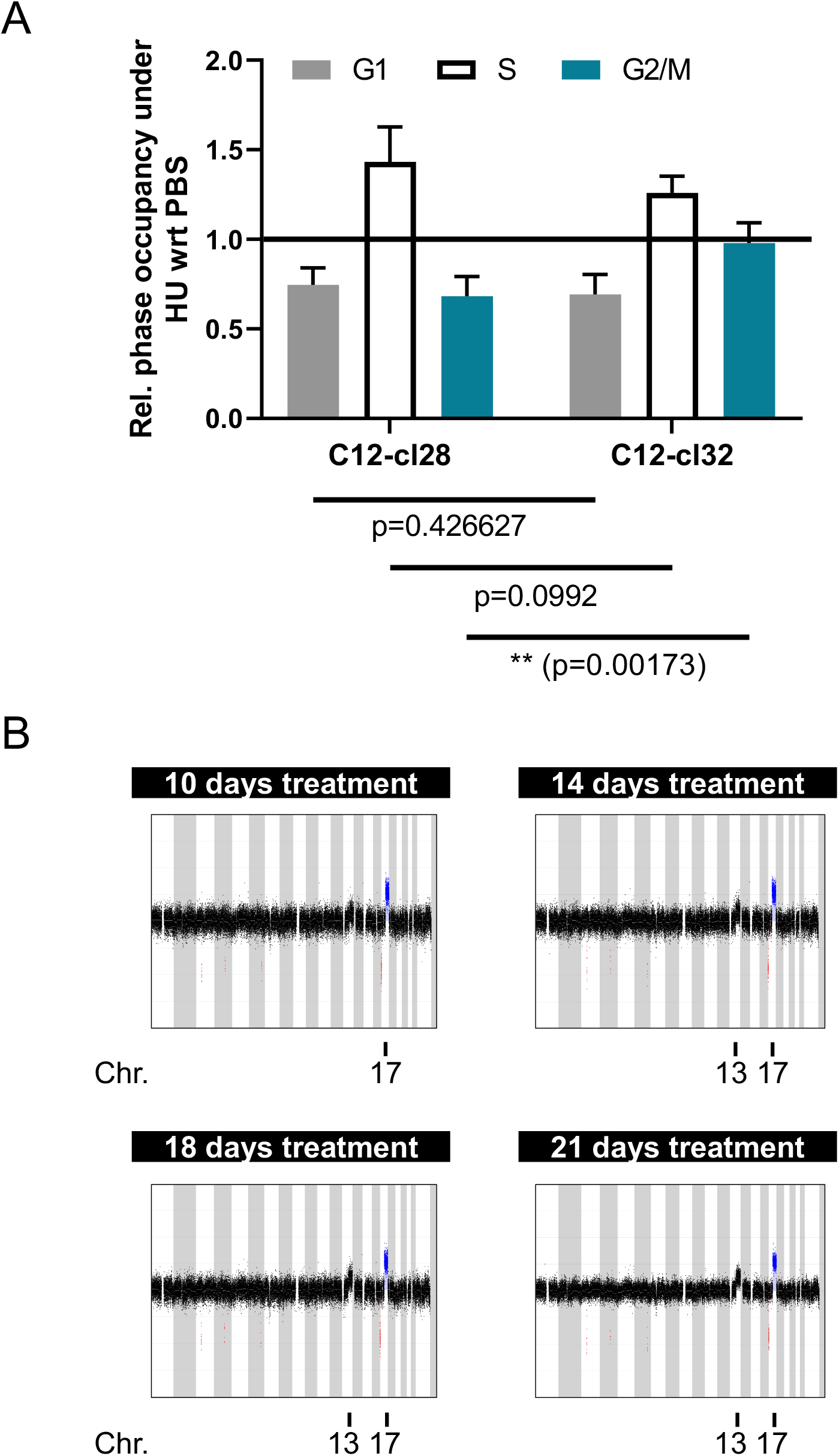
Acquired chromosomal imbalances including 17q gain attenuate hydroxyurea-induced G1/S-phase arrest. **[A]** Flow cytometry-based cell cycle analysis of C12-cl28 and - cl32 under control conditions (PBS) and replicative stress (150μM HU). Bar plot represents relative phase occupancy of cells under replicative stress with regards to control conditions (n=4 biological replicates (d10, d14, d18, d21), see also **[B]** and **Fig. S2**); mean with error bars representing SD after error propagation) **[B]** Matching chromosomal profiles of samples included in the cell cycle analysis by sWGS of C12-cl32 cultured under continuous replicative stress (150μM HU). (*p<0.05; **p<0.01; ***p<0.001)

To further assess the effects of the presence of 17q+ in hESCs, we measured proliferation rates of C12-cl28 and -cl32 cells under normal versus HU conditions. Cumulative growth analyses using the 3T3 assay (**Fig. 3A)** demonstrated comparable proliferation rates of C12-cl28 and -cl32 cells under normal conditions. HU-treated C12-cl32 cells however, showed a statistically significantly increased growth rate (**Fig. 3A** **& S2C-D** p-value < 0.001), in keeping with our sWGS and ddPCR results.

**Figure 3:**
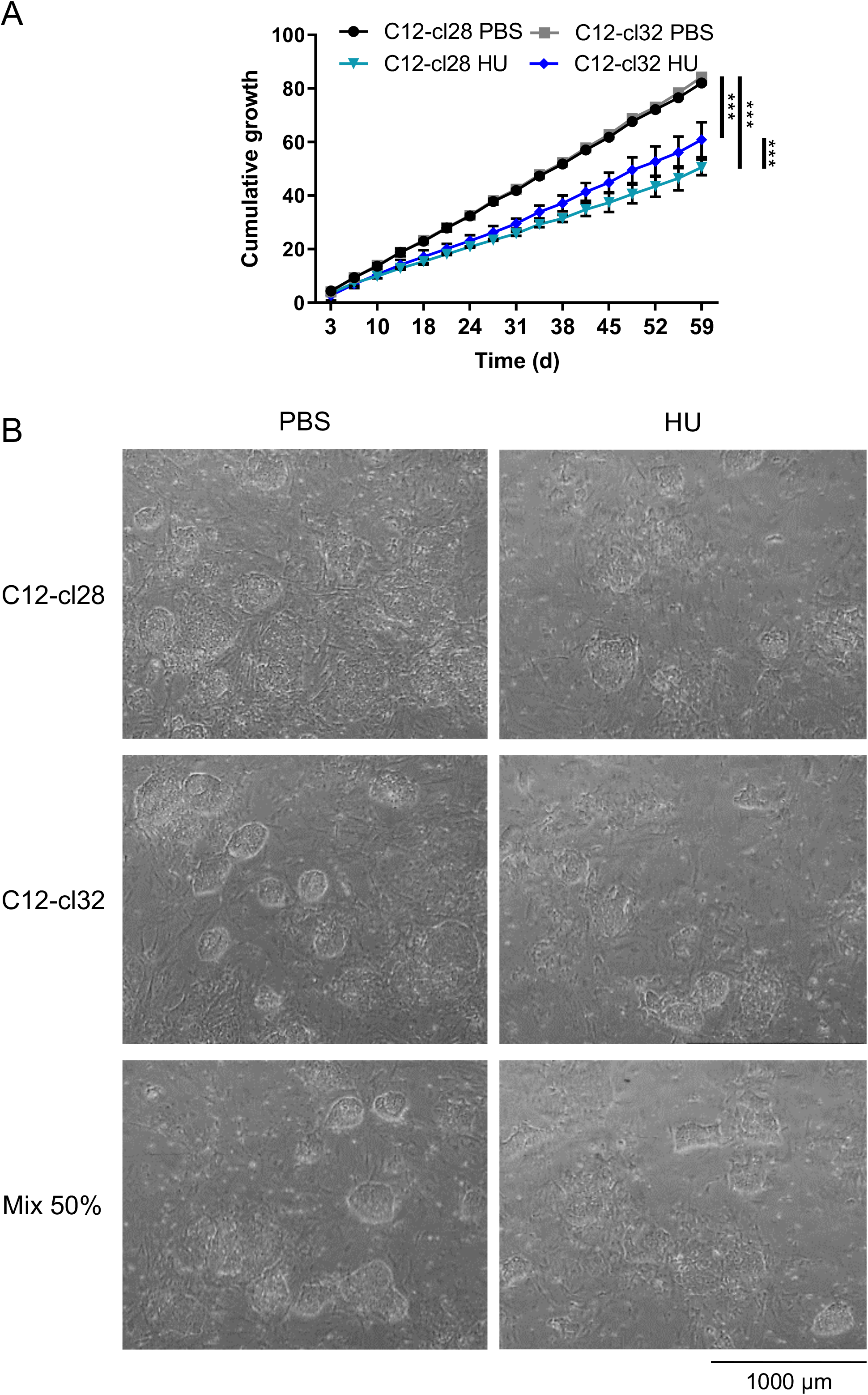
Acquired chromosomal imbalances including 17q gain provide a proliferative advantage. **[A]** Cumulative growth of C12-cl28 and -cl32 under control conditions (PBS) and replicative stress (150μM HU) over time (days). (n=2 biological replicates; mean with error bars representing SD after error propagation, see also **Fig. S2**) [**B]** Morphology of naïve cultures during mixing experiment, grown on MEF feeder layer, as assessed by phase contrast imaging under control conditions (PBS) and replicative stress (150μM HU; day 14). (*p<0.05; **p<0.01; ***p<0.001)

Furthermore, microscopic observations of cultured hESCs during the mixing experiment revealed that the normal dome-shaped colony morphology of tightly packed cells (**Fig. 3B**) altered towards colonies with loosely packed cells and loss of the dome-shape appearance under replicative stress for both clones. These cells display an enlarged appearance and adopt a flattened morphology which suggests a more differentiated/primed-like phenotype under conditions of replicative stress.

### Comparative transcriptome analysis of normal versus 17q+ hESCs under standard and enhanced replicative stress growth conditions

To gain insight into gene dosage effects driving the above described phenotypic effects of 17q+ of HU-treated hESCs, we performed transcriptome analysis of the clones during the early phases of the co-culture experiment (passage 1-6; in total 48 samples; n=12 biological replicates). Gene set enrichment analysis (GSEA) analysis using the C2 MSigDB collection for the expression profiles of C12-cl28 versus C12-cl32 hESCs under HU, revealed significant enrichment for several gene sets (“NIKOLSKY BREAST CANCER 17Q21_25 AMPLICON” and “LASTOWSKA NEUROBLASTOMA COPY NUMBER UP”; control treatment data not shown, HU data see **Fig. 4A**), related to 17q copy number changes in C12-cl32 cells, supporting the validity of our analyses. Of further interest, ZEB1 repressed targets were found among the top upregulated gene sets of our transcriptome analysis for HU-treated C12-cl32 hESCs (**Fig. 4A**).

**Figure 4:**
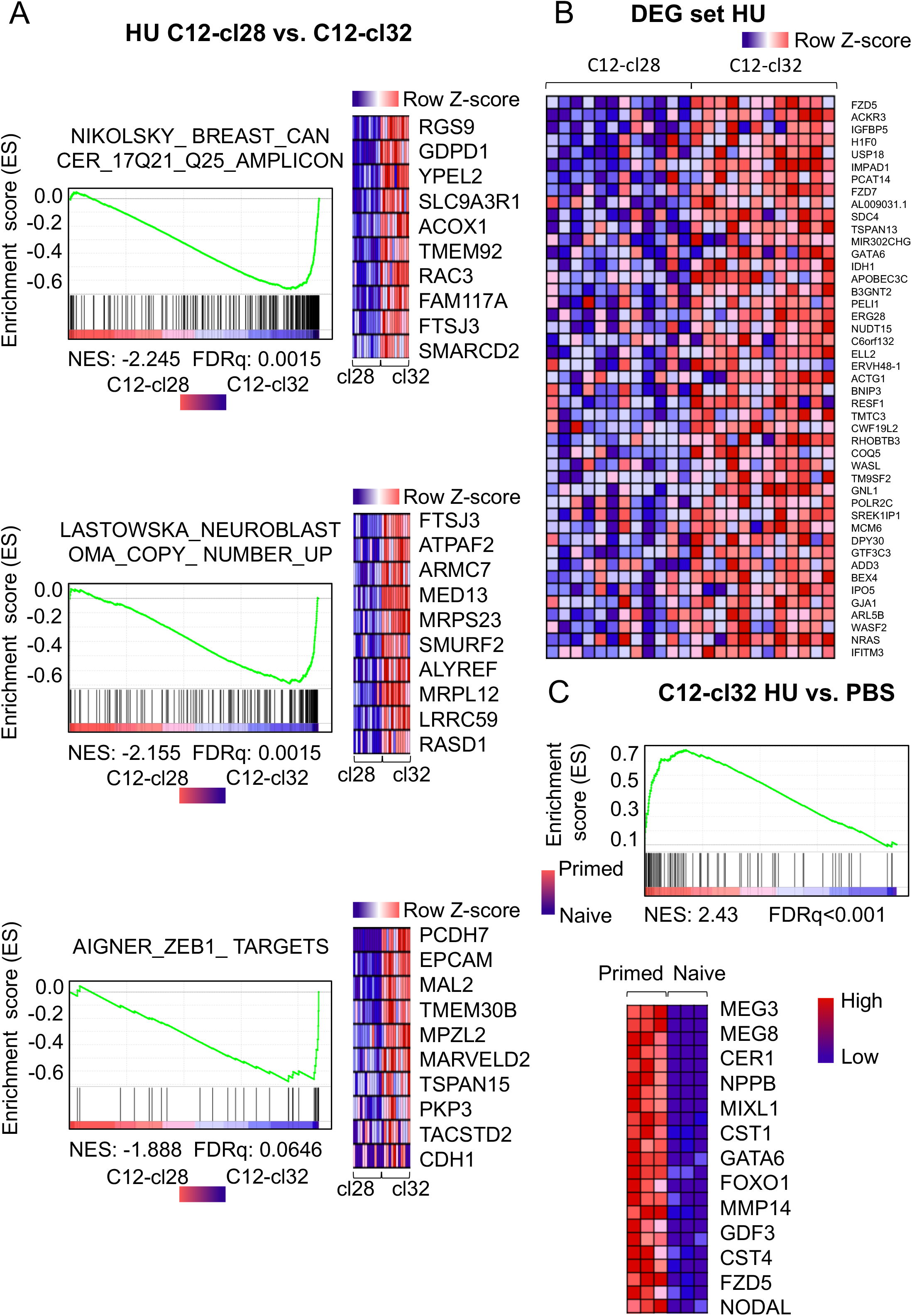
Comparative transcriptome analysis of normal versus 17q+ hESCs under standard and replicative stress imposed growth conditions. (n=12: 2 biological replicates with each 6 passages of independent cultures) **[A]** Top enriched upregulated genesets for C12-cl32 compared to C12-cl28 under HU exposure (150μM) as identified by GSEA. The according heatmap displays the top-10 genes of the core enrichment. (tested for 21 720 genes) **[B]** Heatmap of significantly upregulated genes in C12-cl32 under HU versus PBS and that are not significantly differentially expressed in C12-cl28 under HU versus PBS. (DEG: differentially expressed genes) **[C]** Genes significantly upregulated in C12-cl32 in conditions of replicative stress (150μM HU) versus normal growing conditions (PBS; tested for 22 920 genes) are significantly enriched in the genes upregulated in primed versus naive hESCs. The according heatmap displays the top genes of the core enrichment.

In a next step, as preliminary effort towards exploring the putative genes driving 17q gain selection, we performed differential gene expression analysis for C12-cl28 versus C12-cl32 cells under HU and excluded genes that were also differentially expressed between the two cell populations under normal conditions (PBS) (**Fig. 4B)**. Gene ontology analysis (amp.pharm.mssm.edu/Enrichr/) for the set of genes significantly upregulated in C12-cl32 versus Cl2-c28 under HU includes, amongst others, components of the WNT pathway (including *FZD5* and *FZD7*) and cytoskeleton organisation (including *WASL* and *ACTG1*) ^11–13^.

Additionally, we evaluated the enrichment of genes differentially expressed in C12-cl28 or C12-cl32 in normal growth conditions versus HU exposure in expression data of primed versus naive hESCs ^8^. We found that only genes upregulated in C12-cl32 under HU are significantly enriched in the gene set associated with primed pluripotency (**Fig. 4C**), which is in concordance with the observed morphological changes.

Finally, we also sought for significant enrichment of gene sets corresponding to specific cytogenetic bands using the C1 MSigDB collection (gsea-msigdb.org). We observed a significant enrichment for 13q genes (**Fig. S3**) in C12-cl32 versus C12-cl28 upon HU treatment, including *ZIC2*, a well-known pluripotency marker, and *SOX21*, implicated in maintenance of progenitors and inhibition of neuronal outgrowth ^14^.

## DISCUSSION

Culture adaptation of hESCs has been attributed to several processes including enhanced cloning efficiency, alteration in the balance between differentiation and self-renewal^15^, enrichment of the pluripotent cell population^16^, growth factor and niche independence^16^, enhanced proliferation^17^ and enrichment of cells in S phase of the cell cycle^18^. The occurrence of recurrent whole chromosome 12 and 17 or segmental 12p and 17q gains in long term cultured hESCs is highly intriguing and largely unexplained. The link between these chromosomal imbalances and cancer has been previously made in relation to testicular germ cell tumors which typically exhibit isochromosome 12p or whole chromosome 12 gains^19^ and neuroblastoma in which whole chromosome 17 and 17q gain are the most frequently occurring imbalances^20^. It is generally assumed that combined dosage effects for multiple genes enhance the fitness of the selected clones. Both the nature of these genes and the involved cellular processes remain however largely unknown. We recently identified TBX2, located on 17q, as member of a core regulatory circuit of transcription factors in adrenergic neuroblastoma cells and proposed that TBX2 enhances MYCN sustained activation of FOXM1 targets, the latter which include multiple cell cycle activated genes implicated in control of DNA replication, DNA damage signalling and replicative stress^21^. In addition, BRCA1, also located on 17q and highly expressed in neuroblastoma cells, was shown to be involved in preventing MYCN-dependent accumulation of stalled RNAPII which indirectly also contributes to avoiding transcription-replication conflicts and subsequent replication fork stalling and replicative stress^22^. Given these observations, we hypothesized that replicative stress may act as a major driver in the emergence of hESC subclones with chromosomal imbalances. To test this, we evaluated the effects of hydroxyurea^10^, a well-known inducer of replicative stress, on the growth properties of a parental hESC lines and an isogenic 17q gain containing subclone and obtained evidence for a proliferative advantage of 17q+ hESCs versus the parental line. While several experiments repeatedly revealed emergence of the 17q+ subclone, remarkably also partial 13q gain (encompassing the MYCN driven oncogenic miR-17~92 cluster) co-emerged with 17q gain under conditions of replicative stress. Next, we showed that the 17q+ hESC subclone showed an attenuated increase in S-phase arrest under HU treatment compared to the euploid parental hESCs, resulting in an increased G2/M-phase cell population. In a first step towards identifying the genes affected by dosage effects driving the above described phenotypic effects of HU-treated 17q+ hESCs, our transcriptome analysis showed an enrichment for 17q copy number related data sets thus validating our analysis but further studies are warranted to unequivocally identify the key targets. Moreover, as we have currently identified several candidate dosage sensitive 17q genes in neuroblastoma (Vanhauwaert et al., unpublished), it will be of interest to compare culprit 17q replicative stress resistors in hESCs and neuroblastoma. This may be of critical importance in the light of ongoing efforts to generate hESC- and iPSC-derived genetically well-defined tumor model ^23,24^. In addition to available iPSC-derived cell lines from neuroblastoma patients with specific driver mutations in amongst others the ALK oncogene (Van Haver et al., unpublished), we also aim to generate hESC or iPSC neuroblastoma models with chromosome 17q gain to study, for the first time, the impact of 17q gene dosage effects on MYCN-driven tumor formation.

In conclusion, we provide the first experimental evidence for a positive selective effect of copy number imbalances such as 17q+ on hESC proliferation under elevated replicative stress culture conditions. Further molecular studies are warranted to dissect the molecular basis of these copy number changes which may also shed light onto their role in cancer cells, with putative implication in identification of novel non-mutated druggable dependencies.

## DATA AVAILABILITY STATEMENT

The data that support the findings of this study are publicly available at GEO (GSE150647).

## ACKNOWLEDGEMENTS

The authors thank F. Matthijssens, R. De Smedt and I. Velghe for the technical support with regards to the FACS experiments and K. Verniers for the technical support with regards to the ddPCR experiments. Further, we would also like to thank the CNVseq team of the diagnostic lab of the Centre of Medical Genetics, Ghent, for their technical support with the sWGS profiling. E. De Meester, S. De Keulenaer and L. Tilleman of NXTGNT are thanked for their support in the transcriptome profiling.

---

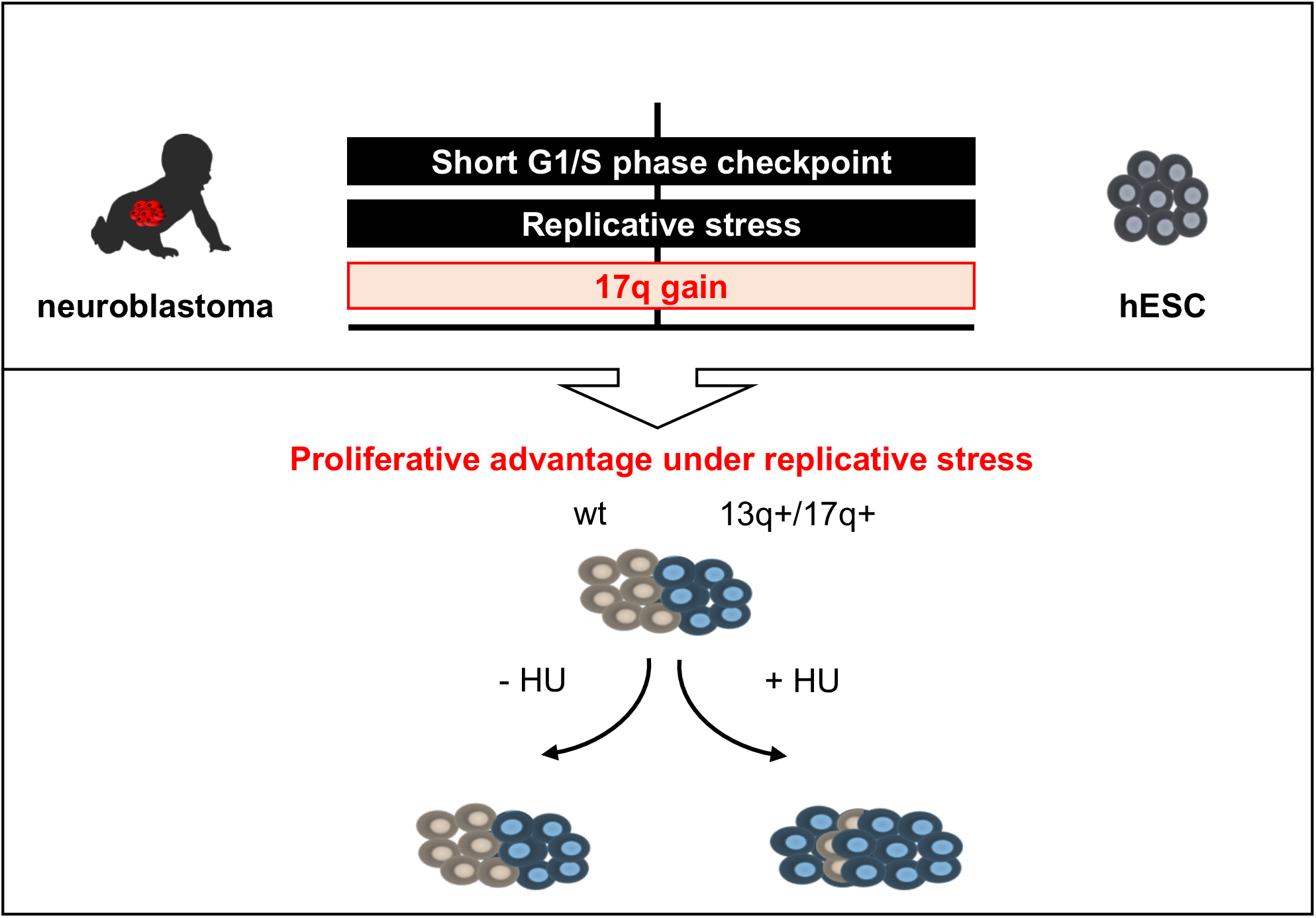

