## Supplemental info for "Recurrent chromosomal imbalances provide selective advantage to human embryonic stem cells under enhanced replicative stress conditions"

**Mus *et. al.***

**Supplemental Table S1, related to figure 1 and supplemental figure S1: Overview of chromosomal aberrations above Vivar calling threshold in late passage C12 (passage 124) and during the mixing experiment in the mix and C12-cl32**

| | C12 (passage 124) | C12-cl32 (passage 54, start mixing experiment) | Mix (42 days HU) | C12-cl32 (42 days PBS)[a] | C12-cl32 (42 days HU)[b] |
|---|---|---|---|---|---|
| gain | 1q25.3q32.1 (20.85 Mb) | 5q35.1q35.1 (0.15 Mb) | 1p36.13p34.2 (17.75 Mb) | 5q35.1q35.1 (0.15 Mb) | 3q28q28 (0.7 Mb) |
| | 3p12.3p12.3 (1.4 Mb) | 17q21.31q25.3 (38.85 Mb) | 5q35.1q35.1 (0.15 Mb) | 17q21.31q25.3 (39.85 Mb) | 5q35.1q35.1 (0.15 Mb) |
| | 7p22.2p11.2 (54.75 Mb) | | 7q11.22q11.22 (1.2 Mb) | | 13q21.32q34 (47.75 Mb) |
| | 7q11.21q36.3 (89 Mb) | | 13q21.32q34 (47.45 Mb) | | 17q21.31q25.3 (39.85 Mb) |
| | 11q23.3q23.3 (0.6 Mb) | | 17q21.31q25.3 (39.85 Mb) | | |
| | 17q21.3q25.3 (39.85 Mb) | | | | |
| | 20q12q12 (0.25 Mb) | | | | |

[a] gain in chromosome 13 and loss in chromosome 18 not included, as value below Vivar calling threshold

[b] gain in chromosome 2 not included, as value below Vivar calling threshold

**Supplemental Table S1, related to figure 1 and supplemental figure S1: continued.**

| | C12 (passage 124) | C12-cl32 (passage 54, start mixing experiment) | Mix (42 days HU) | C12-cl32 (42 days PBS)[a] | C12-cl32 (42 days HU)[b] |
|---|---|---|---|---|---|
| loss | 8q11.1q11.1 (0.7 Mb) | 3p14.2p14.2 (0.65 Mb) | 3p14.2p14.2 (0.6 Mb) | 3p14.2p14.2 (0.6 Mb) | 3p14.2p14.2 (0.65 Mb) |
| | 16p13.3p13.3 (0.7 Mb) | 4q25q26 (0.35 Mb) | 4q25q26 (0.35 Mb) | 4q25q26 (0.4 Mb) | 4q25q26 (0.35 Mb) |
| | 19p13.2p13.2 (0.5 Mb) | 6q23.3q23.3 (0.3 Mb) | 6q23.3q23.3 (0.3 Mb) | 6q23.3q23.3 (0.3 Mb) | 6q23.3q23.3 (0.3 Mb) |
| | 19q13.41q13.41 (0.8 Mb) | 17p13.3p13.2 (3.3 Mb) | 17p13.3p13.2 (3.3 Mb) | 17p13.3p13.2 (3.3 Mb) | 17p13.3p13.2 (3.3 Mb) |
| | X 23p11.3 (0.65 Mb) | | | | X 23p11.3 (0.2 Mb) |
| | X 23p11.23p11.23 (1.05 Mb) | | | | |
| | X 23q22.3q22.3 (0.25 Mb) | | | | |

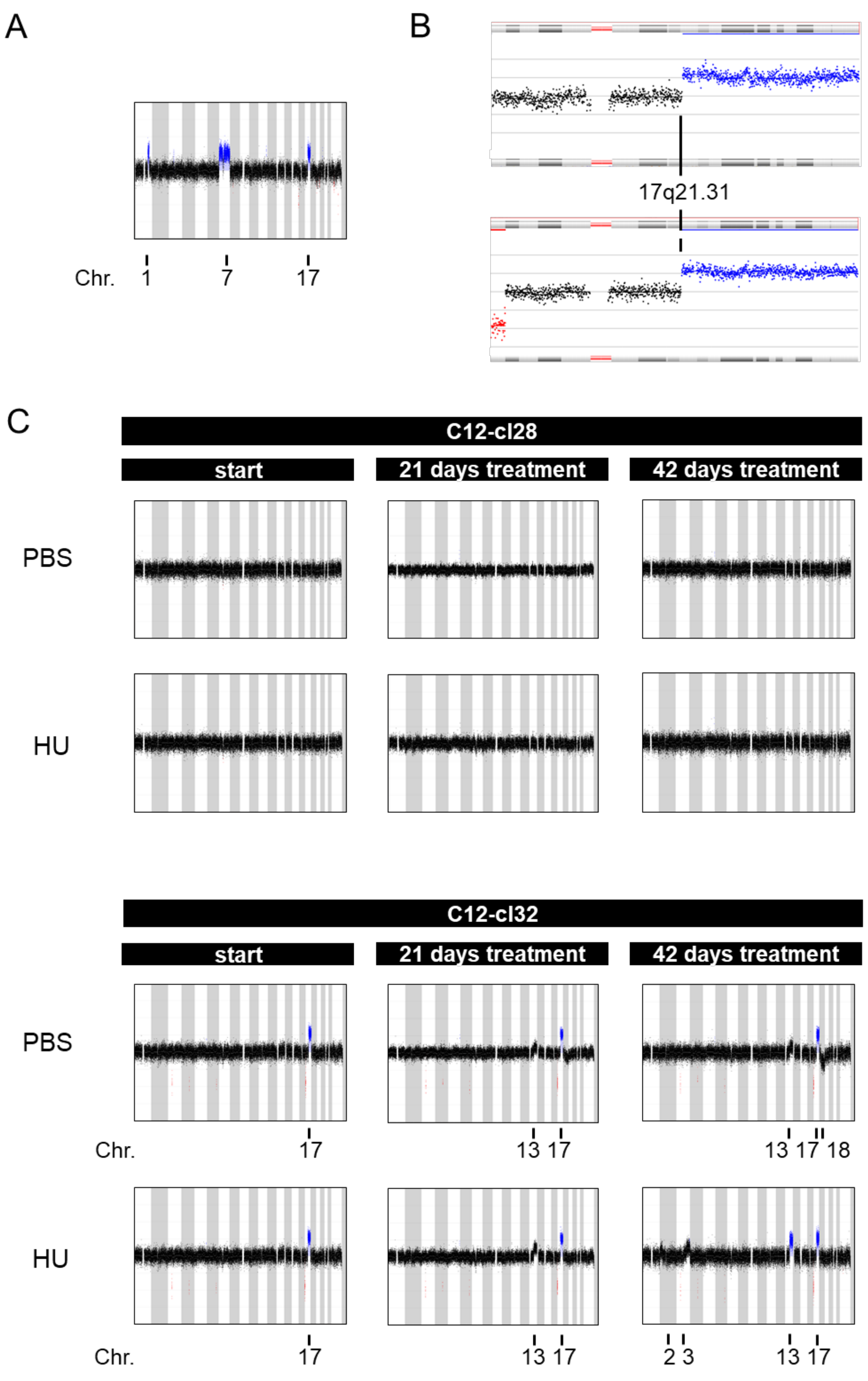


**Supplemental Figure S1, related to figure 1: Recurrent acquired chromosomal imbalances confer a proliferative advantage to hESCs under conditions of replicative stress.** **[A]** Chromosomal profiles by sWGS of C12 with a gain in chromosome 1q25.3q32.1, 7 and 17q21.31q25.3 (passage 124) **[B]** sWGS line view on chromosome 17 breakpoint of gain for the cells in panel **[A]** (upper panel, passage 124) and C12-cl32 (lower panel, start mixing experiment, passage 54) **[C]** Chromosomal profiles by sWGS of C12-cl28 and C12-cl32 over time (days), cultured under control conditions (PBS) and RS (150µM HU). (representative of n=2 shown (biological replicates))

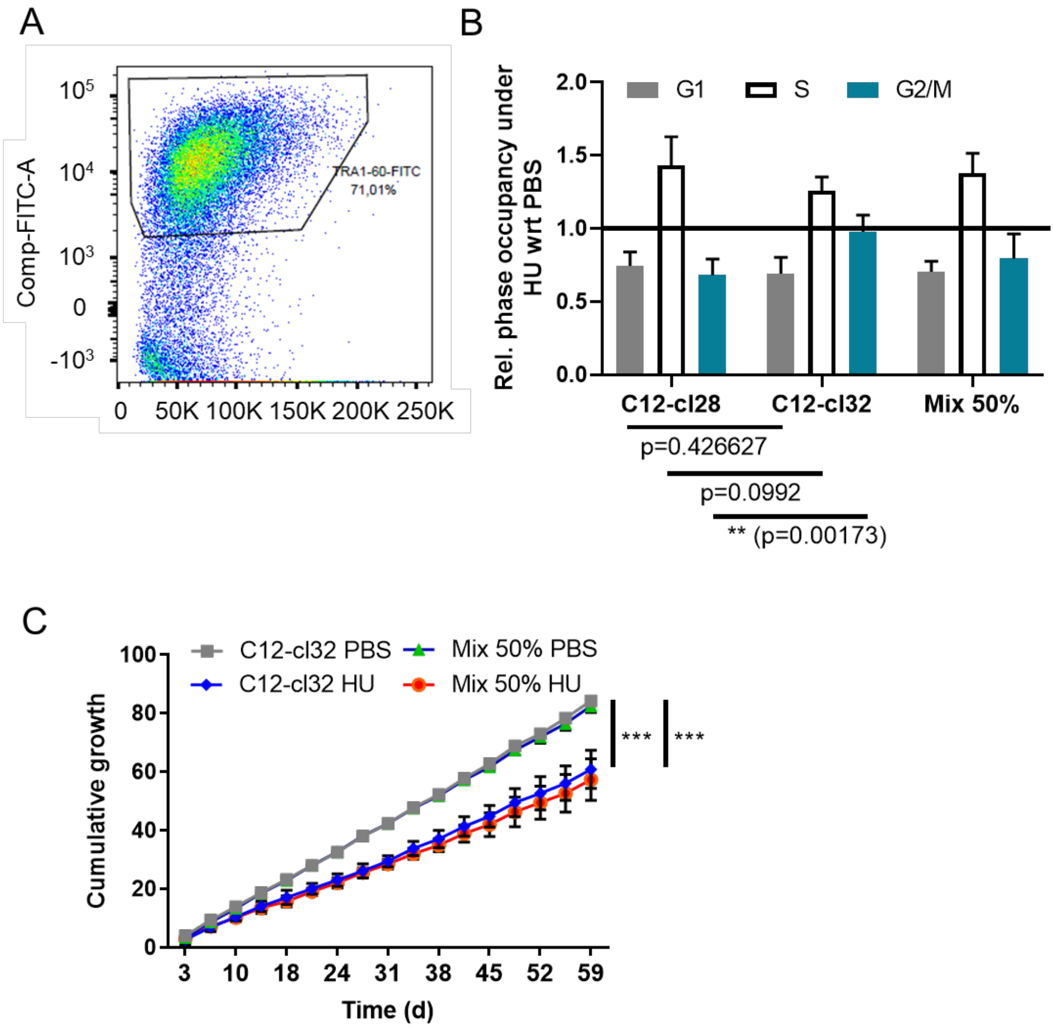


D

| t.test equal slope | C12_cl28_PBS | C12_cl32_PBS | mix_PBS | C12_cl28_HU | C12_cl32_HU | mix_HU |
|---|---|---|---|---|---|---|
| C12_cl28_PBS | | | | | | |
| C12_cl32_PBS | 0.7136 | | | | | |
| mix_PBS | 0.7984 | 1 | | | | |
| C12_cl28_HU | < 0.0001 | < 0.0001 | < 0.0001 | | | |
| C12_cl32_HU | < 0.0001 | < 0.0001 | < 0.0001 | < 0.0001 | | |
| mix_HU | < 0.0001 | < 0.0001 | < 0.0001 | < 0.0001 | 0.998 | |

**Supplemental Figure S2, related to figure 2-3: Acquired chromosomal imbalances including 17q gain attenuate hydroxyurea induced G1/S-phase arrest and provide a proliferative advantage. [A]** The C12-cl28 hESC under control conditions are separated from the MEF feeder layer based on a TRA-1-60-FITC hESC specific marker during FACS analysis (n=4 biological replicates; representative figure shown for all conditions and cell lines of **Fig. 2**). **[B]** Flow cytometry based cell cycle analysis of C12-cl28 and C12-cl32 and co-culture (mix 50%) under control conditions (PBS) and RS (150µM HU). Bar plot represent relative phase occupancy of cells under RS with regards to control conditions (n=4 biological replicates; mean with error bars representing SD after error propagation) **[C]** Cumulative growth of C12-cl32 and the co-culture (mix 50%) under control conditions (PBS) and RS (150 µM HU) over time (days). (n=2 biological replicates; mean with error bars representing SD after error propagation) **[D]** Overview of pairwise test for equal slope (cumulative growth) between C12-cl28, C12-cl32 and the co-culture under control conditions (PBS) and under continuous RS (150 µM HU). (n=2 biological replicates)

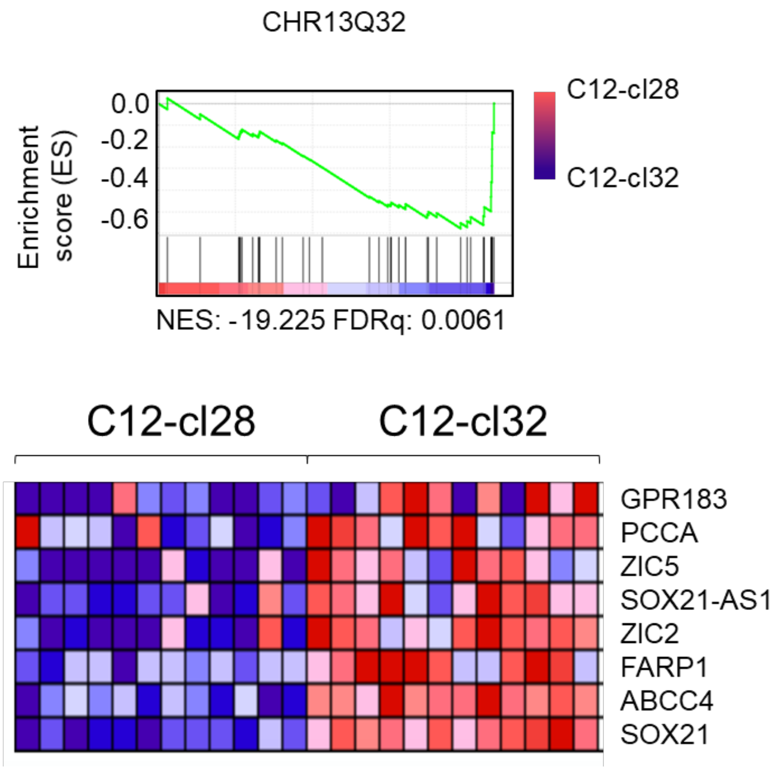


**Supplemental Figure S3, related to figure 4: 13q gain enrichment in C12-cl32 upon replicative stress.** Enrichment of chromosome 13q32 gene signature as identified using GSEA for C12-cl28 vs. C12-cl32 under conditions of RS (150µM HU; tested for 21 720 genes), the leading edge gene list is given in the lower panel.
